# Discovery of Selective Nrf2 Activators from Natural Products: A Computational Screening Approach to Minimize Off-Target Effects on PXR and CYP2D6

**DOI:** 10.64898/2026.04.12.718057

**Authors:** Yujin Wang, Yiwen Gong, Ruofei Li, Zhiru Li, Hongxia Cai, Li Fan, Yutong Li, Haomiao Ma

## Abstract

Nuclear factor erythroid 2-related factor 2 (Nrf2) is a central regulator of cellular antioxidant responses and a highly promising therapeutic target for a range of oxidative stress-related diseases. However, the clinical translation of Nrf2 activators has been hampered by significant off-target effects—notably unintended activation of the pregnane X receptor (PXR) and inhibition of cytochrome P450 2D6 (CYP2D6)—which can lead to dangerous drug-drug interactions and metabolic complications. To overcome this critical barrier, we conducted the first large-scale computational screening of 628,898 natural products from the COCONUT database, integrating molecular docking with a rigorous three-tier selectivity strategy designed to prioritize compounds that strongly bind KEAP1 (the primary Nrf2 repressor) while minimizing interactions with PXR and CYP2D6. Our innovative approach identified 10 ultraselective candidates that demonstrate potent KEAP1 affinity, negligible PXR engagement, and only moderate CYP2D6 binding—achieving up to 12.29-fold selectivity for Nrf2 pathway activation. These top hits are structurally novel, enriched in lipid-like and nucleoside-inspired scaffolds, and exhibit promising drug-like properties. By providing both a curated set of chemically diverse, selectivity-optimized leads and a publicly accessible screening dataset, this work establishes a new foundation for the rational development of safer, more precise Nrf2-targeted therapies, bridging a crucial gap between target potential and clinical viability. By prioritizing compounds with minimal off-target effects on PXR and CYP2D6, our approach offers a scalable template for reducing drug development failures and advancing safer therapeutics for oxidative stress-related diseases.

Graphical abstract

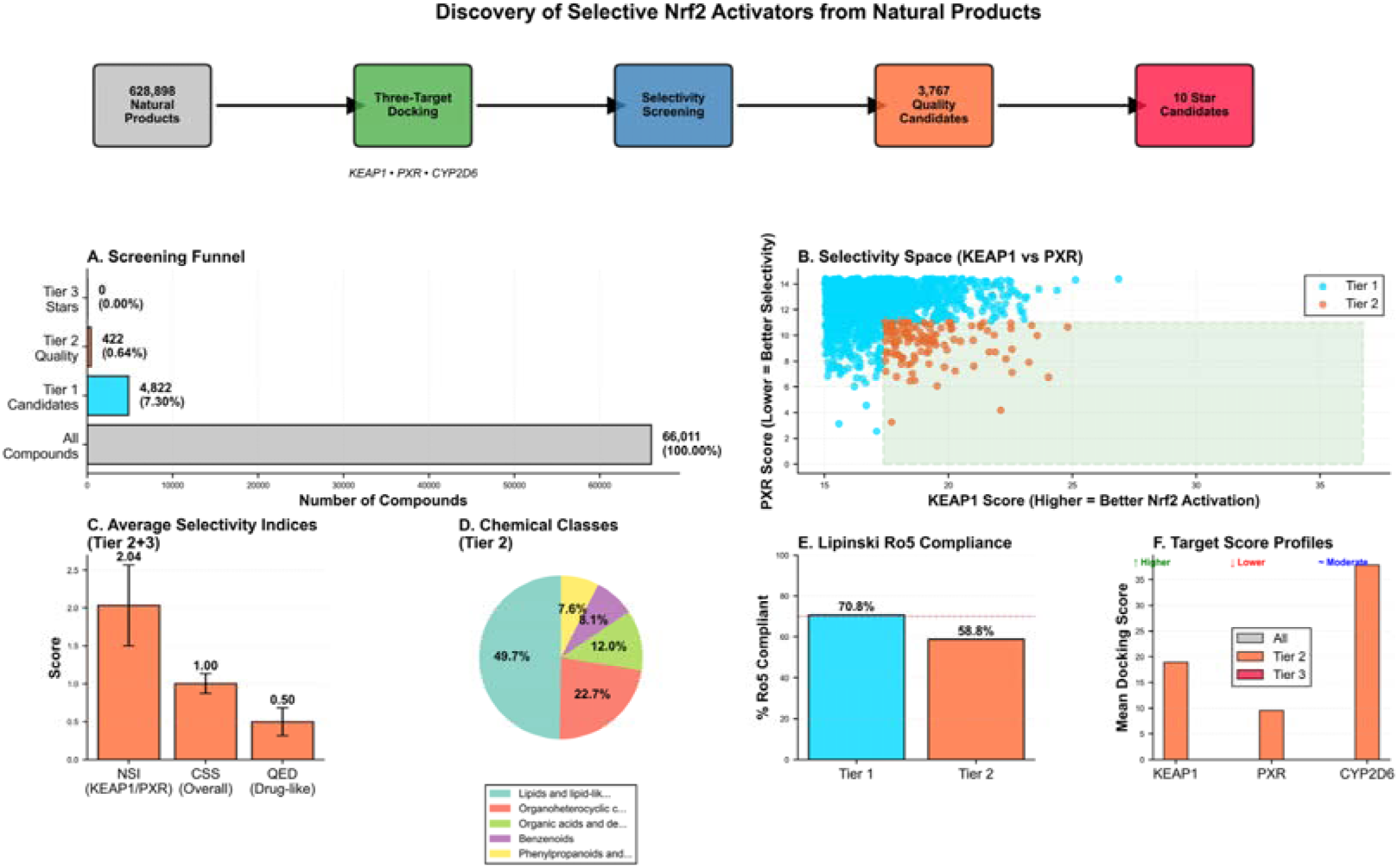

## 1. Introduction

The nuclear factor erythroid 2-related factor 2 (Nrf2) signaling pathway is one of the most critical cellular defense systems against oxidative stress, electrophilic damage, and inflammation. [1–3] By orchestrating the transcription of over 250 cytoprotective genes, [3,4] Nrf2 governs redox homeostasis, detoxification, and metabolic adaptation. Its pharmacological activation has therefore emerged as a compelling therapeutic strategy for a wide spectrum of diseases, including neurodegenerative disorders, cardiovascular disease, chronic kidney disease, and cancer chemoprevention. [4–6] Despite its therapeutic promise, the translation of Nrf2 activation into clinically safe drugs has been repeatedly hindered by off-target toxicities. [6,7] In particular, many small-molecule activators inadvertently activate the pregnane X receptor (PXR)—a master regulator of xenobiotic metabolism—leading to severe drug-drug interactions (DDIs), [7,8] while others potently inhibit cytochrome P450 2D6 (CYP2D6), disrupting the metabolism of approximately a quarter of all prescribed drugs. [9,10] These dual off-target liabilities have imposed significant barriers to clinical application, emphasizing the need for selective Nrf2 activators that balance efficacy with pharmacological safety.

Natural products provide an unparalleled chemical reservoir for discovering safer bioactive molecules. Shaped by evolutionary selection, they often exhibit high target specificity and three-dimensional complexity that synthetic libraries rarely achieve. [11,12] Indeed, several natural compounds, such as sulforaphane, curcumin, and resveratrol, have shown the ability to activate Nrf2 and confer cytoprotective effects. [13–15] Yet, existing studies have focused primarily on potency rather than selectivity, and systematic efforts to identify natural Nrf2 activators devoid of PXR or CYP2D6 liabilities remain scarce. Addressing this gap requires not only large-scale exploration of natural product chemical space but also a rigorous evaluation of off-target interactions within the same screening framework.

To address these gaps, our study aims to bridge target potential and clinical viability by implementing a safety-by-design framework. Here, we report a large-scale computational investigation of natural products for selective Nrf2 activation conducted to date. Leveraging the COCONUT database, [16] encompassing 628,898 structurally diverse natural products, we implemented a three-target molecular docking strategy against KEAP1 (the negative regulator of Nrf2), PXR, and CYP2D6. To systematically evaluate therapeutic potential, we designed a three-tier selectivity screening pipeline and introduced quantitative indices—the Nrf2 Selectivity Index (NSI), Safety Index (SI), and Comprehensive Selectivity Score (CSS)—to prioritize compounds that strongly engage KEAP1 while minimizing interactions with PXR and CYP2D6. This integrative approach enables simultaneous assessment of efficacy and toxicological safety at an unprecedented scale.

Our analysis yielded tens of thousands of selective hits and uncovered novel structural motifs with remarkable selectivity profiles. Notably, lipid-based scaffolds and nucleoside analogues were found to be significantly enriched among high-performing candidates, suggesting unexplored chemical territories for safe Nrf2 modulation. Beyond identifying promising leads, this study generates a comprehensive selectivity-focused dataset—a unique open resource that bridges molecular docking, toxicological filtering, and cheminformatic profiling.

Taken together, this work introduces a large-scale, data-driven pipline for uncovering natural product scaffolds that selectively activate Nrf2 while minimizing off-target risks, laying the foundation for safer therapeutic development guided by early toxicological insights.

## 2. Materials and Methods

### 2.1 Data Sources and Integration

#### 2.1.1 COCONUT Natural Product Database

The natural product library was obtained from the COCONUT database (https://coconut.naturalproducts.net), which aggregates structural information from over 50 natural product resources. [16] We downloaded version 2023.11 containing 696,707 unique natural products with associated chemical structures (SMILES format), molecular descriptors, and biological source annotations. Each compound entry includes standardized identifiers, molecular properties (molecular weight, LogP, hydrogen bond donors/acceptors, topological polar surface area), drug-likeness metrics (QED score, Lipinski violations), structural features (aromatic ring count, rotatable bonds, fraction of sp³ carbons), and chemical classifications (superclass, class, NP Classifier pathway annotations).

#### 2.1.2 Docking Score Data

Molecular docking calculations were performed using the KARMA (Kinetic And Rigid-body Molecular docking with Autodock) scoring function [17] against three protein targets:

KEAP1 (Kelch domain): Target for Nrf2 activation, using PDB structure [23] representing the binding site for Nrf2; PXR (Pregnane X Receptor): Ligand-binding domain to assess DDI risk; CYP2D6 (Cytochrome P450 2D6): Active site to evaluate metabolic inhibition potential

Docking was performed for 628,898 natural products that had corresponding entries in the COCONUT database. For each target, three scoring metrics were generated: (1) karma_score (primary docking score), (2) karma_score_ff (force field-based score), and (3) karma_score_aligned (alignment-corrected score). The karma_score was used as the primary metric for all subsequent analyses, as it represents the most reliable estimate of binding affinity. Higher scores indicate stronger predicted binding interactions.

#### 2.1.3 Data Integration Pipeline

The three docking datasets (KEAP1, PXR, CYP2D6) were merged using compound identifiers as the common key, resulting in a unified dataset containing 628,898 compounds with docking scores across all three targets. This merged dataset was then joined with the COCONUT database using a left join to preserve all molecular property information. The final integrated dataset contained 38 variables including docking scores, molecular descriptors, drug-likeness parameters, and chemical classification annotations.

### 2.2 Selectivity Screening Strategy

#### 2.2.1 Rationale for Three-Tier Screening

To systematically identify compounds with optimal selectivity profiles, we developed a three-tier screening funnel with progressively stringent criteria This approach allows for the identification of a broad candidate pool (Tier 1) for exploratory analyses, a refined set of quality candidates (Tier 2) for prioritization, and an elite group of star compounds (Tier 3) for immediate experimental validation. The rationale for integrating PXR and CYP2D6 into our screening strategy stems from their well-established roles in drug-induced toxicity. PXR activation is a known culprit in drug-drug interactions (DDIs) due to its regulation of xenobiotic metabolism, [7] while CYP2D6 inhibition can disrupt the metabolism of ∼25% of prescribed drugs, leading to adverse effects. [9] By including these targets, we adopt a safety-by-design principle that preemptively addresses toxicological risks. This approach contrasts with traditional methods that often prioritize potency alone, resulting in late-stage failures. Our three-tier funnel thus serves as a proactive tool for embedding toxicological safety into the hit identification process.

#### 2.2.2 Screening Criteria

Percentile thresholds were calculated for each target’s docking scores to define objective cutoffs:

Tier 1 (Candidate Pool) - Relaxed Criteria: KEAP1 score > 50th percentile (>15.01), PXR score < 50th percentile (<14.44) CYP2D6 score between 30th-70th percentile (30.36-43.74)

Tier 2 (Quality Candidates) - Stringent Criteria: KEAP1 score > 70th percentile (>17.40), PXR score < 30th percentile (<11.08) CYP2D6 score between 30th-70th percentile (30.36-43.74)

Tier 3 (Star Candidates) - Very Stringent Criteria: KEAP1 score > 90th percentile (>20.90), PXR score < 10th percentile (<6.26) CYP2D6 score between 40th-60th percentile (33.71-39.77)

The rationale for these cutoffs reflects the therapeutic objectives: strong KEAP1 binding is desired for potent Nrf2 activation; minimal PXR binding reduces DDI risk; moderate CYP2D6 binding (avoiding extremes) minimizes metabolic perturbations while maintaining acceptable safety margins.

It is worth noting that the 40–60th percentile range for CYP2D6 binding was intentionally selected to represent moderate interaction rather than complete absence of binding. Extremely weak CYP2D6 affinity may indicate poor overall metabolic recognition or unrealistic docking behavior, whereas moderate engagement reflects compounds that are likely to be metabolically processed without acting as strong inhibitors. This middle range therefore provides a practical balance, capturing molecules with acceptable metabolic compatibility while avoiding those that could cause excessive inhibition or poor clearance.

### 2.3 Selectivity Index Development

#### 2.3.1 Nrf2 Selectivity Index (NSI)

NSI quantifies the preferential binding to KEAP1 over PXR:

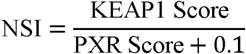

The constant 0.1 prevents division by zero for compounds with minimal PXR binding. Higher NSI values indicate greater selectivity for KEAP1, with NSI > 2 considered favorable.

#### 2.3.2 Safety Index (SI)

SI assesses the balance between KEAP1 activity and CYP2D6 inhibition:

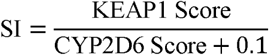

Higher SI values suggest stronger KEAP1 binding relative to CYP2D6 inhibition. However, excessively high SI may indicate very weak CYP2D6 interactions, which could paradoxically suggest poor overall binding affinity.

#### 2.3.3 Comprehensive Selectivity Score (CSS)

CSS integrates both selectivity dimensions into a single metric:

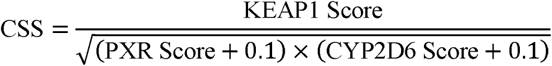

CSS provides a balanced assessment of overall selectivity, rewarding strong KEAP1 binding while penalizing high PXR or CYP2D6 scores. CSS was used as the primary ranking criterion for identifying top candidates.

### 2.4 Drug-Likeness and ADME Property Analysis

#### 2.4.1 Lipinski’s Rule of Five

Compliance with Lipinski’s Rule of Five [18] was assessed using pre-calculated violations in the COCONUT database. Compounds were classified as fully compliant (0 violations), acceptable (1 violation), or poor (≥2 violations). Individual parameters were analyzed including: Molecular weight ≤ 500 Da; ALogP ≤ 5; Hydrogen bond donors ≤ 5 and Hydrogen bond acceptors ≤ 10.

#### 2.4.2 QED Drug-Likeness

Quantitative Estimate of Drug-likeness (QED) scores, which integrate multiple molecular properties into a unitless metric (0-1 scale), [19] were obtained from the COCONUT database. QED ≥ 0.5 was considered favorable for oral drug development.

#### 2.4.3 Additional ADME-Relevant Properties

In addition to standard drug-likeness criteria, several ADME-relevant physicochemical properties were evaluated to further characterize compound developability. The topological polar surface area (TPSA) was examined as a key determinant of oral absorption, with an optimal range of 20–140 Å² for balancing permeability and solubility. [20] Rotatable bond count was used as an indicator of molecular flexibility, where values ≤10 were preferred to enhance membrane permeability and conformational stability. The fraction of sp³-hybridized carbons (Fsp³) was included as a proxy for three-dimensionality, as higher Fsp³ values have been associated with improved safety profiles and reduced promiscuous binding. [21] Finally, aromatic ring count was analyzed because lower aromaticity generally correlated with greater selectivity in preliminary analyses, suggesting that compounds with fewer aromatic rings may exhibit reduced off-target interactions. [22]

### 2.5 Chemical Classification and Enrichment Analysis

#### 2.5.1 Chemical Class Annotations

Chemical classifications for all compounds were obtained from the COCONUT database annotations to facilitate structural and biosynthetic analyses. Each molecule was categorized across multiple hierarchical levels, including chemical superclass (broad structural category), chemical class (intermediate structural grouping), and chemical subclass (specific structural family). In addition, NP Classifier annotations were incorporated to capture biosynthetic origins, including pathway, superclass, and class levels, thereby linking chemical structure to potential natural product biosynthetic lineages.

#### 2.5.2 Enrichment Analysis

For each tier, chemical class distributions were compared to the background distribution (all screened compounds) to identify enriched or depleted classes. Enrichment factor was calculated as:

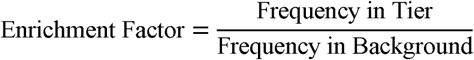

Enrichment factors > 1.5 were considered significant over-representation.

### 2.6 Statistical and Multivariate Analyses

#### 2.6.1 Correlation Analysis

Spearman rank correlation coefficients were calculated between all molecular properties, docking scores, and selectivity indices using SciPy (v1.11). [25] Correlations with |ρ| > 0.3 were considered moderate, and |ρ| > 0.5 were considered strong.

#### 2.6.2 Principal Component Analysis

PCA was performed on 14 standardized molecular features (docking scores and physicochemical properties) using scikit-learn (v1.3). [24] Features were standardized using StandardScaler before PCA to ensure equal weighting. Principal components explaining >5% variance were retained for interpretation.

#### 2.6.3 Statistical Comparisons

Mann-Whitney U tests were used to compare continuous variables between tiers due to non-normal distributions. Statistical significance was set at p < 0.05 (two-tailed). [25]

### 2.7 Data Availability

All integrated data, screening results, and analysis code are available in the project repository at https://doi.org/10.5281/zenodo.17562589. The COCONUT database is publicly accessible at https://coconut.naturalproducts.net/.

## 3. Results and Discussion

### 3.1 Overview of Docking Score Distributions Across Three Targets

We successfully integrated docking scores for 628,898 natural products across the three critical targets: KEAP1, PXR, and CYP2D6 (Figure 1). The docking score distributions revealed distinct patterns for each target that informed our subsequent selectivity screening strategy. KEAP1 scores exhibited a relatively narrow distribution (mean: 14.92 ± 4.74, range: 0.00-51.89), suggesting that most natural products have moderate predicted binding affinity to the KEAP1 Kelch domain. This distribution is consistent with the notion that KEAP1 binding requires specific structural features to engage the shallow, relatively flat binding pocket. [1,2]

**Figure 1.**
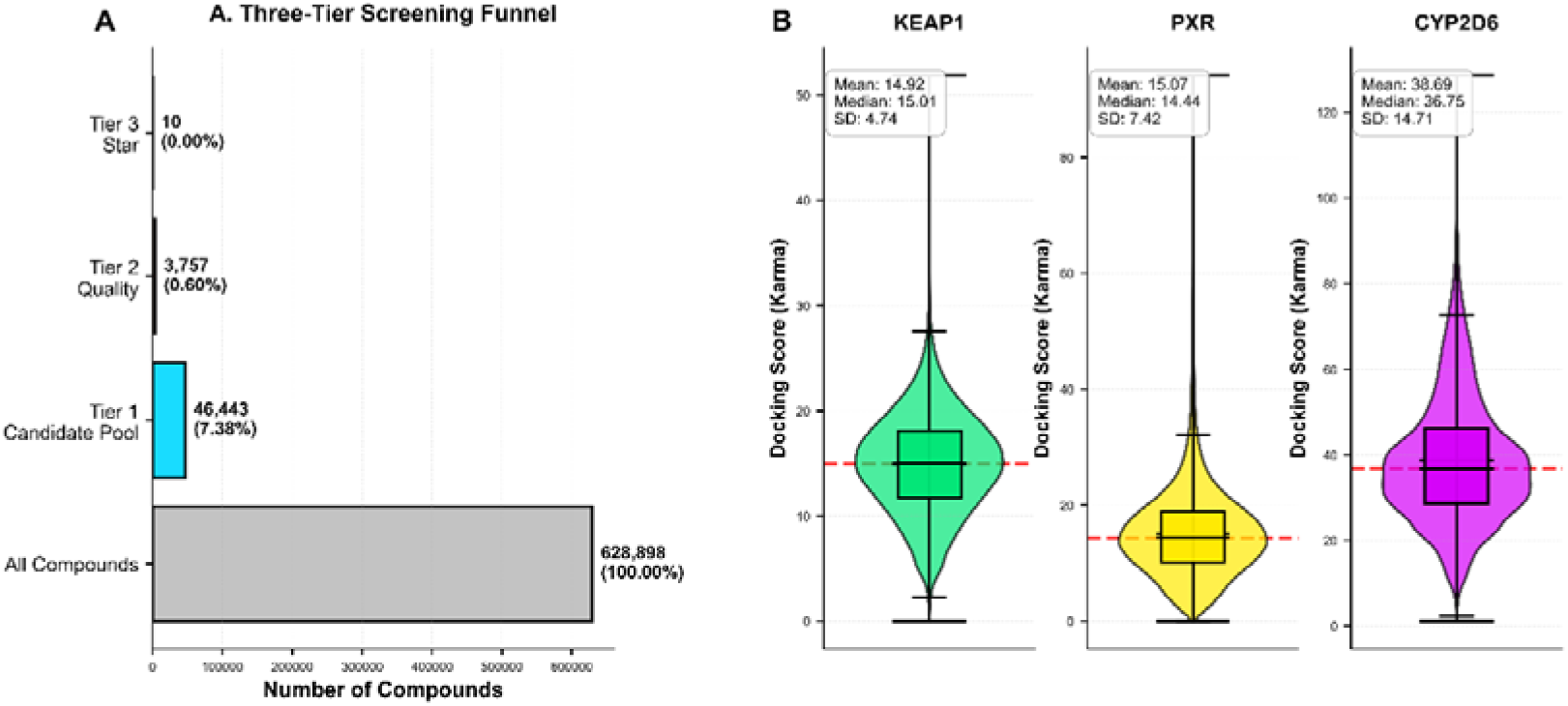
Three-Tier Screening Funnel for Selective Nrf2 Activators. **(A)** The screening cascade applied progressively stringent selectivity criteria to identify compounds with optimal KEAP1 binding, minimal PXR activation, and moderate CYP2D6 interaction profiles. Starting from 628,898 natural products, we identified 46,443 Tier 1 candidates (7.38%), 3,767 Tier 2 quality candidates (0.60%), and 10 ultra-selective Tier 3 star candidates (0.002%). The dramatic reduction in compound numbers with increasing stringency demonstrates the rarity of natural products with exceptional selectivity profiles. **(B)** Docking Score Distributions Across Three Targets. Violin plots with overlaid box plots show the distribution of docking scores for KEAP1, PXR, and CYP2D6 across all 628,898 screened natural products. Red dashed lines indicate percentile thresholds (10th, 30th, 50th, 70th, 90th) used for tier definitions.

In contrast, PXR scores showed a broader distribution (mean: 15.07 ± 7.42, range: 0.00-94.18,) with greater variance, reflecting the well-known promiscuity of the PXR ligand-binding domain. [7,8] The larger ligand-binding pocket of PXR is known to accommodate diverse chemical structures, which explains the wider range of predicted binding affinities observed in our dataset. This promiscuity is precisely what makes PXR activation a concern for drug development, as even structurally diverse compounds can inadvertently activate this nuclear receptor. [8]

CYP2D6 displayed the highest mean docking scores (mean: 38.69 ± 14.71, range: 1.13-128.70) with the greatest variability. The higher absolute scores likely reflect differences in the scoring function calibration rather than fundamentally stronger binding, as CYP2D6 has a relatively constrained active site. [9,10] The large standard deviation suggests substantial chemical diversity in how natural products interact with the CYP2D6 active site, with some compounds showing very strong predicted inhibition while others show minimal interaction.

The three-tier screening strategy successfully enriched for compounds with progressively improved selectivity profiles (Figure 1, Table 1). Tier 1 (Candidate Pool) identified 46,443 compounds (7.38% of total), representing a substantial yet manageable set for furtheranalysis. These compounds met relaxed criteria requiring above-median KEAP1 binding, below-median PXR binding, and moderate CYP2D6 interactions. The relatively high pass rate for Tier 1 reflects the fact that approximately 25% of compounds would randomly satisfy these criteria if the three scores were independent; the observed 7.38% pass rate indicates substantial correlation between the target scores.

**Table 1.** Summary Statistics of Screening Tiers.

| Tier | Count | % of | KEAP1 | PXR (mean | CYP2D6 | NSI (mean | SI (mean± | CSS (mean |
| --- | --- | --- | --- | --- | --- | --- | --- | --- |
|  |  | Total | (mean±SD) | ±SD) | (mean±SD) | ±SD) | SD) | ±SD) |
| All | 628,898 | 100.00% | 14.92±4.74 | 15.07±7.42 | 38.69±14.71 | 1.15±1.45 | 0.40±0.10 | 0.66±0.19 |
| Tier 1 | 46,443 | 7.38% | 17.38±2.24 | 10.25±2.23 | 37.25±4.09 | 1.75±0.53 | 0.47±0.06 | 0.81±0.10 |
| Tier 2 | 3,767 | 0.60% | 19.28±1.91 | 8.28±2.06 | 37.38±4.03 | 2.48±0.78 | 0.52±0.05 | 1.01±0.14 |
| Tier 3 | 10 | 0.002% | 22.45±1.34 | 4.22±1.54 | 37.62±2.32 | 5.78±2.77 | 0.60±0.03 | 1.68±0.37 |

The distinct docking score distributions for KEAP1, PXR, and CYP2D6 reveal fundamental insights into target promiscuity and safety risks. The narrow distribution of KEAP1 scores suggests a selective binding pocket, whereas the broad PXR distribution underscores its well-known promiscuity, which often leads to DDIs in clinical settings. [7,8] Similarly, the high variability in CYP2D6 scores highlights the metabolic diversity of natural products. These patterns justify our safety-by-design approach, where stringent criteria for PXR and CYP2D6 are necessary to preemptively filter out compounds with high off-target potential. The three-tier funnel effectively enriches for compounds that balance efficacy and safety, as visualized in Figure 1.

Tier 2 (Quality Candidates) identified 3,767 compounds (0.60% of total), applying more stringent criteria that required strong KEAP1 binding (>70th percentile), minimal PXR binding (<30th percentile), and moderate CYP2D6 binding. The Tier 2 compounds represent our primary pool of quality candidates suitable for experimental validation and further development.

Tier 3 (Star Candidates) identified only 10 compounds (0.002% of total), requiring exceptional KEAP1 binding (>90th percentile), negligible PXR binding (<10th percentile), and tightly controlled CYP2D6 binding (40th-60th percentile). The extreme rarity of Tier 3 compounds, that representing only 1 in 60,000 natural products highlights the exquisite selectivity challenge (Figure 2). These ultraselective compounds warrant immediate prioritization for experimental validation.

**Figure 2.**
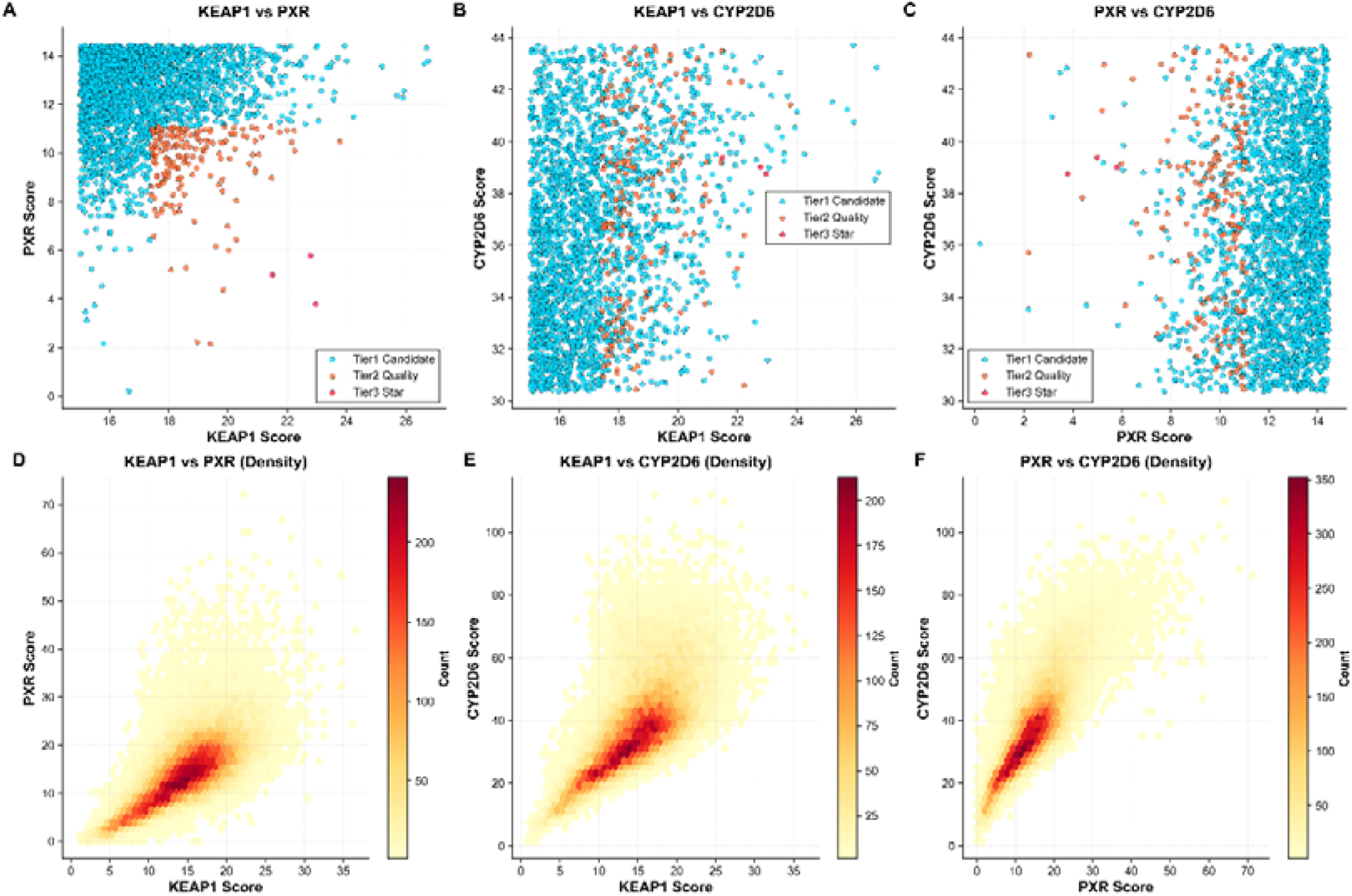
Pairwise Target Score Relationships. Scatter plots showing the relationship between docking scores for **(A)** KEAP1 vs PXR, **(B)** KEAP1 vs CYP2D6, and **(C)** PXR vs CYP2D6. **(D-F)** Corresponding hexbin density plots showing the concentration of compounds in 2D docking score space. The plots reveal that selective compounds (Tier 2 and 3) occupy distinct regions of chemical space characterized by high KEAP1, low PXR, and moderate CYP2D6 scores.

### 3.2 Selectivity Index Development and Validation

To quantitatively assess selectivity beyond simple score thresholds, we developed three complementary selectivity indices: Nrf2 Selectivity Index (NSI), Safety Index (SI), and Comprehensive Selectivity Score (CSS). These metrics provide continuous measures of selectivity that can be used for ranking and prioritization.

NSI measures the ratio of KEAP1 to PXR binding, with higher values indicating greater preference for KEAP1 (Figure 3A,3D). Across all compounds, NSI showed a median of 1.06, indicating that most natural products bind KEAP1 and PXR with similar affinity. However, Tier 2 compounds achieved mean NSI of 2.48±0.78, representing approximately 2.4-fold selectivity for KEAP1 over PXR. Remarkably, Tier 3 compounds reached mean NSI of 5.78±2.77, with the best compound (CNP0371205.2) achieving NSI of 12.29, indicating >12-fold selectivity for KEAP1. The dramatic increase in NSI across tiers validates this metric as an effective measure of PXR selectivity.

**Figure 3.**
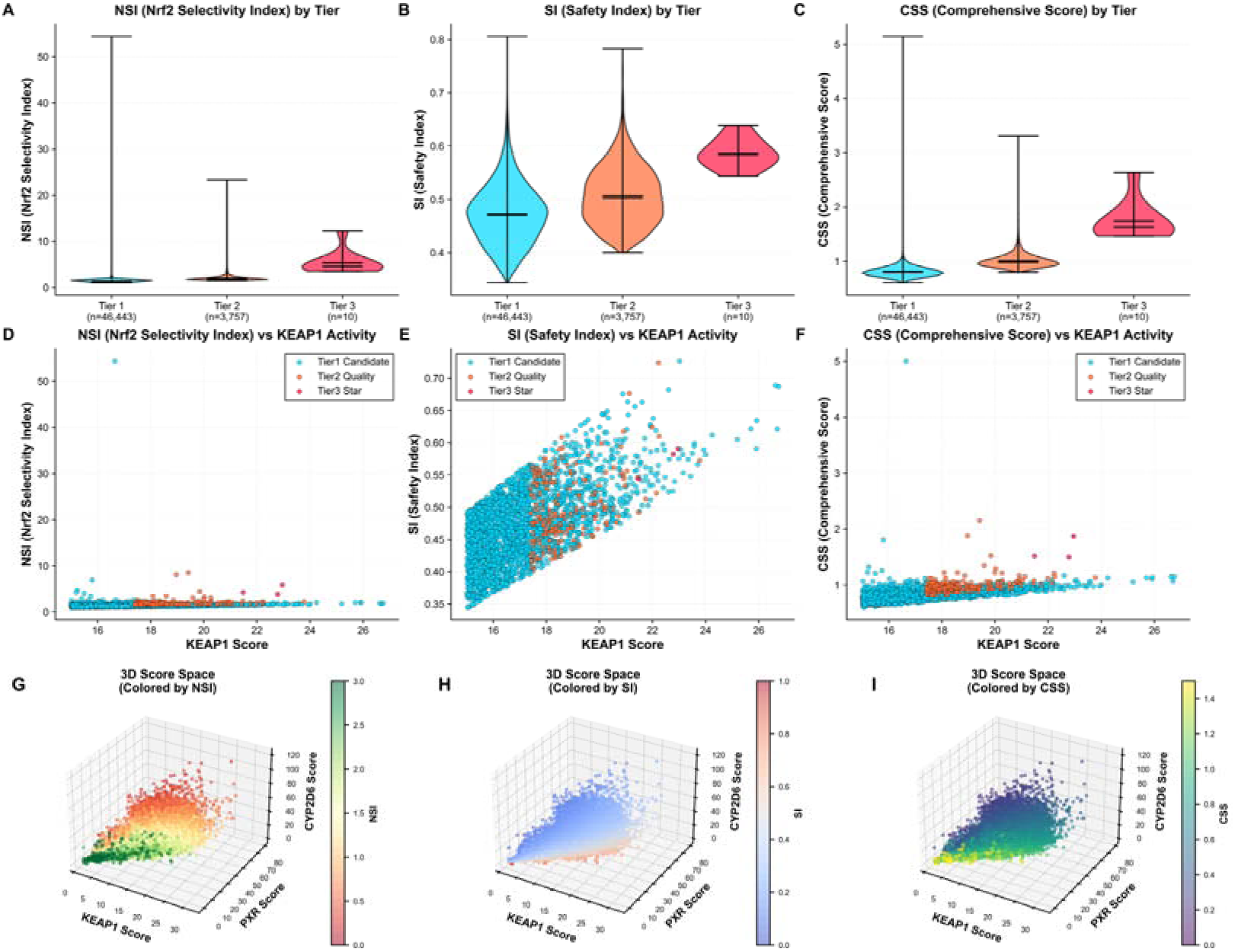
Selectivity Index Distributions and Relationships. **(A-F)** Violin plots showing the distribution of NSI, SI, and CSS across the four tiers. Higher KEAP1 scores generally correlate with improved selectivity indices, particularly for NSI and CSS. The positive relationship between KEAP1 activity and selectivity suggests that potent Nrf2 activation and selectivity are not mutually exclusive goals. **(G-I)** Three views of the same 3D scatter plot showing KEAP1 (x-axis), PXR (y-axis), and CYP2D6 (z-axis) docking scores

SI assesses the balance between KEAP1 activity and CYP2D6 inhibition (Figure 3B,3E). While SI showed more modest variation across tiers (All: 0.40±0.10; Tier 2: 0.52±0.05; Tier 3: 0.60±0.03), the consistent increase demonstrates that selective candidates maintain favorable CYP2D6 profiles. The relatively narrow range of SI values reflects our deliberate strategy to select compounds in the moderate CYP2D6 binding range rather than at extremes.

CSS integrates both selectivity dimensions (PXR and CYP2D6) into a single metric. CSS showed the most dramatic progression across tiers (All: 0.66±0.19; Tier 2: 1.01±0.14; Tier 3: 1.68±0.37), with Tier 3 compounds achieving 2.5-fold higher CSS than the overall population (Figure 3C,3F). We used CSS as the primary ranking metric for identifying top candidates, as it provides a balanced assessment of overall selectivity.

To visualize the multi-dimensional selectivity landscape, we plotted compounds in 3D space defined by KEAP1, PXR, and CYP2D6 scores (Figure 3G-3I). This analysis reveals that selective compounds occupy a distinct region. Tier 3 star candidates cluster tightly in the optimal corner of this space, demonstrating exceptional consistency in their selectivity profiles.

Coloring the 3D space by NSI reveals a clear gradient, with the highest NSI values concentrated in regions of high KEAP1 and low PXR scores, as expected from the NSI definition. Similarly, coloring by CSSshows that compounds with the highest comprehensive selectivity occupy a narrow “sweet spot” in the 3D landscape. The relatively sparse population of this optimal region explains why only 10 compounds achieved Tier 3 status out of nearly 630,000 screened.

The 3D visualization also reveals interesting patterns in the global distribution. Most compounds cluster in a central region with moderate scores across all three targets, reflecting the tendency of natural products to show modest, non-selective binding. A subset of compounds shows high scores across all three targets, representing promiscuous binders that would likely cause significant off-target effects.

The positive correlation between QED and CSS (Spearman ρ=0.430) further suggests that drug-likeness and safety are synergistic, supporting the safety-by-design paradigm. These indices can be adapted to other drug discovery campaigns to quantify off-target risks early, potentially transforming how toxicity is assessed in high-throughput screening.

### 3.3 Drug-Likeness and ADME Property Assessment

A critical consideration for drug development is whether selective candidates possess favorable pharmacokinetic properties. We conducted comprehensive analysis of drug-likeness parameters across all tiers to assess the developability of our selective candidates.

#### 3.3.1 Lipinski’s Rule of Five Compliance

Analysis of Lipinski’s Rule of Five compliance revealed that selective candidates maintain good drug-like properties. Among Tier 1 candidates, 71.2% showed zero Ro5 violations, 23.5% had one violation, and only 5.3% had two or more violation. This high compliance rate is encouraging and significantly better than typical natural product libraries, which often show 40-50% non-compliance. [11] Tier 2 candidates showed slightly reduced compliance (57.8% with zero violations, 33.0% with one violation, 9.2% with ≥2 violations), reflecting a trade-off between selectivity and ideal drug-likeness properties. Interestingly, Tier 3 star candidates showed mixed compliance (40% with zero violations, 40% with one violation, 20% with ≥2 violations), though the small sample size (n=10) limits statistical interpretation.

Individual Lipinski parameters showed distinct patterns (Figure 4). Molecular weight distributions were generally favorable across all tiers, with mean values well below the 500 Da threshold (Tier 1: 382±95 Da; Tier 2: 437±98 Da; Tier 3: 466±72 Da). The progressive increase in molecular weight across tiers suggests that achieving exceptional selectivity may require somewhat larger molecules, possibly to fill selectivity pockets in the KEAP1 Kelch domain (Figure 4A).

**Figure 4.**
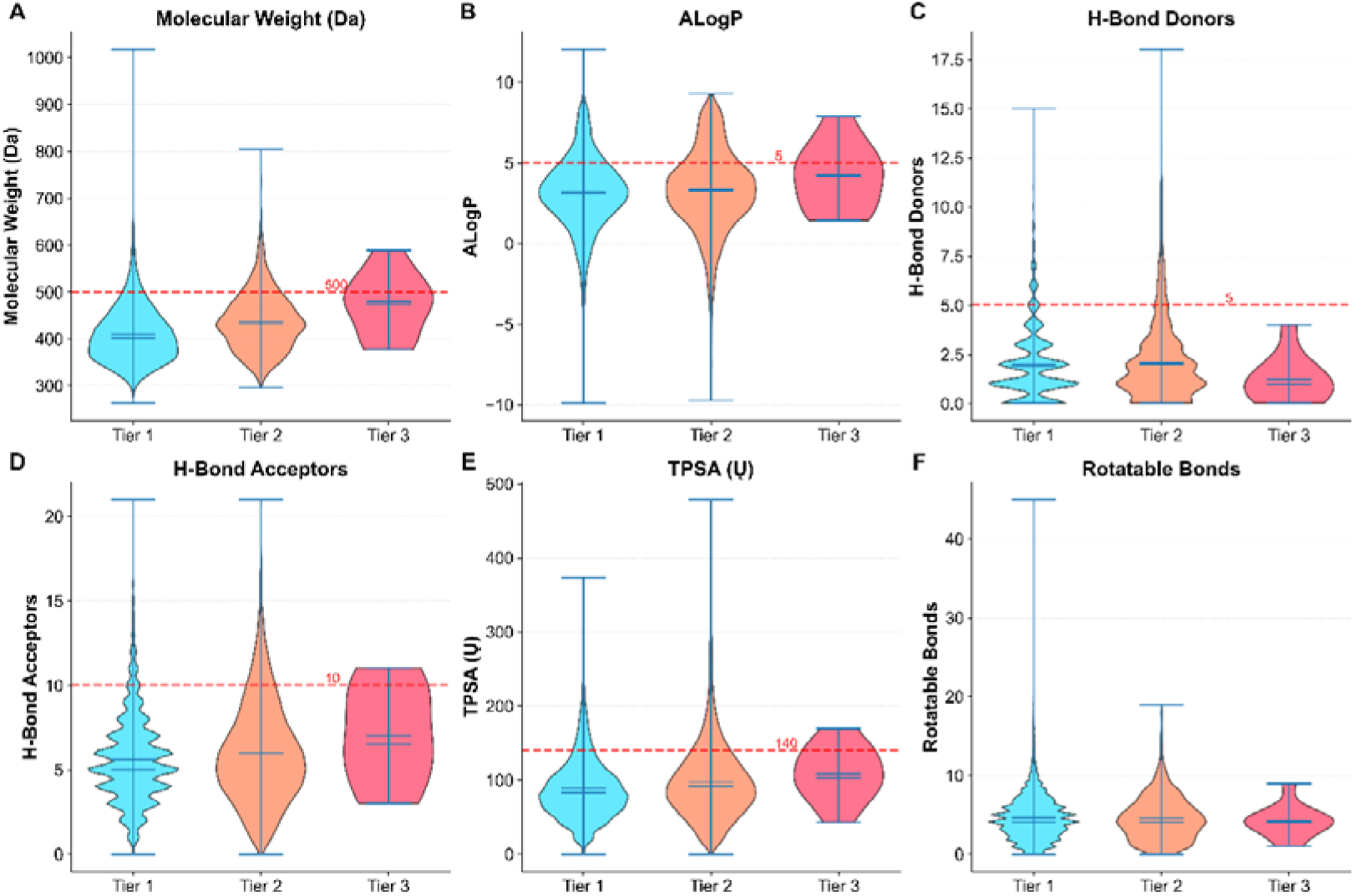
Lipinski Rule of Five Properties Across Tiers. Violin plots showing the distribution of six key drug-likeness parameters across all compounds and three selectivity tiers: **(A)** Molecular weight (Da), **(B)** ALogP, **(C)** Hydrogen bond donors, **(D)** Hydrogen bond acceptors, **(E)** Topological polar surface area (TPSA, Ų), and **(F)** Rotatable bond count. Red dashed lines indicate Lipinski Rule of Five or optimal cutoffs. Despite progressive increase in selectivity from Tier 1 to Tier 3, the compounds maintain generally favorable drug-like properties, with most parameters remaining within acceptable ranges. The slight elevation in molecular weight and LogP for higher tiers suggests that achieving exceptional selectivity may require somewhat larger, more lipophilic scaffolds.

ALogP values showed interesting variations, with Tier 2 candidates exhibiting mean ALogP of 3.64±2.31, comfortably within the Ro5 limit of 5 (Figure 4B). However, a subset of highly selective compounds showed elevated LogP values (up to 9), suggesting that lipophilicity may contribute to KEAP1 binding. Hydrogen bond donors and acceptors remained well within acceptable ranges across all tiers, indicating that selective candidates do not rely on excessive hydrogen bonding for KEAP1 engagement.

TPSA (topological polar surface area) showed mean values around 100-120 Ų across tiers, within the optimal range for oral bioavailability (20-140 Ų, Figure 4E). [20] Rotatable bond counts averaged 7-9 across tiers, acceptable for oral drugs though approaching the upper limit of flexibility. These ADME-relevant properties suggest that selective candidates, particularly from Tier 1 and 2, possess favorable pharmaceutical profiles suitable for further development.

#### 3.3.2 QED Drug-Likeness Assessment

Quantitative Estimate of Drug-likeness (QED) provides an integrated metric of overall drug-likeness on a 0-1 scale [19] (Figure 5). The mean QED score across all natural products was 0.42±0.16, consistent with previous reports that natural products generally have moderate drug-likeness. Interestingly, selective candidates showed comparable QED scores (Tier 1: 0.46±0.15; Tier 2: 0.44±0.15; Tier 3: 0.37±0.14), indicating that selectivity screening does not dramatically compromise drug-likeness.

**Figure 5.**
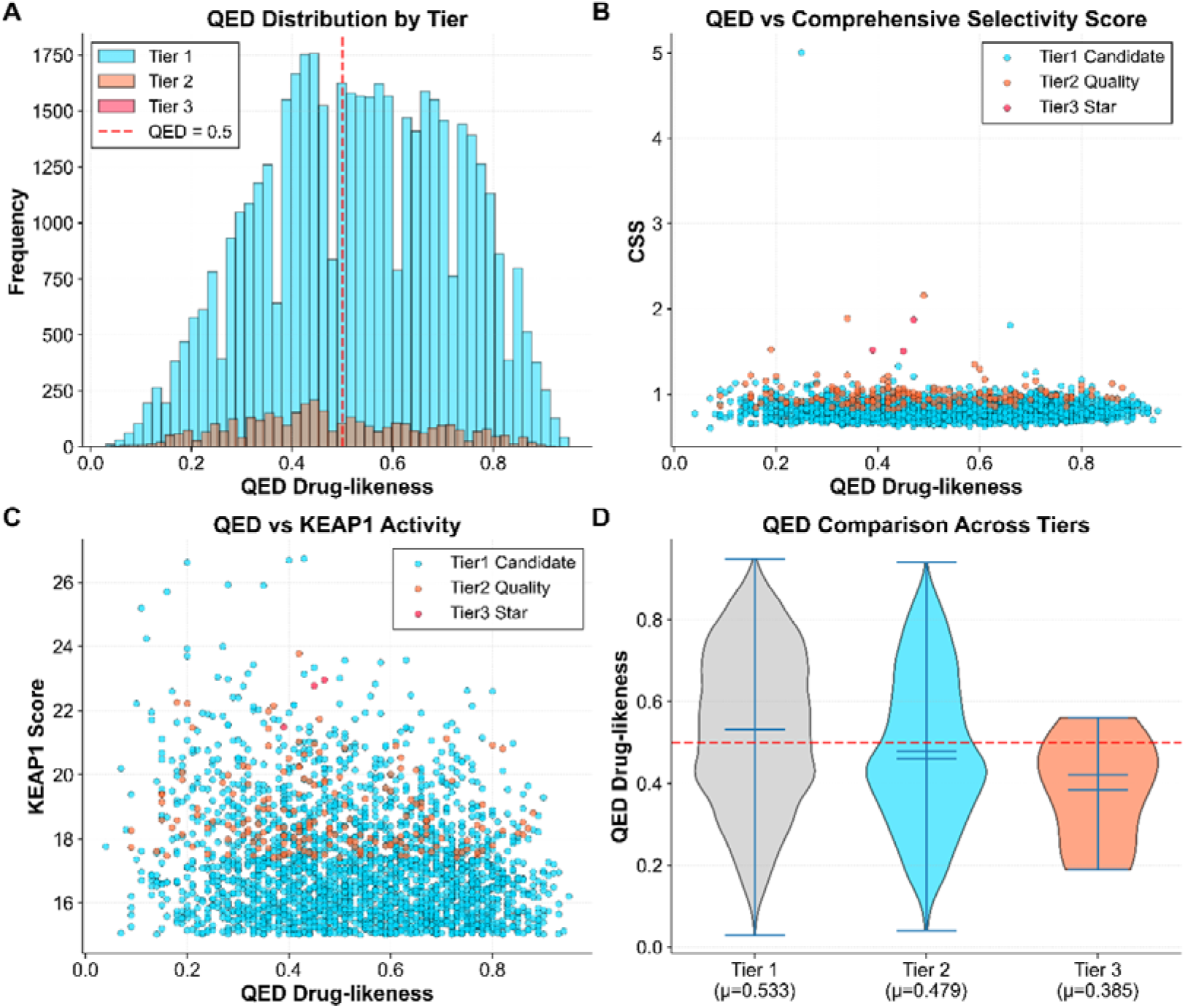
QED Drug-Likeness Analysis. **(A)** Distribution of QED scores across tiers, showing that selective candidates maintain moderate to good drug-likeness despite stringent selectivity criteria. **(B)** Scatter plot of QED versus CSS, revealing a weak positive correlation (ρ=0.430) suggesting compatibility between drug-likeness and selectivity**. (C)** QED versus KEAP1 activity, showing no strong correlation. **(D)** Violin plot comparison of QED distributions, with mean values annotated. Several Tier 2 and 3 compounds achieve the desirable combination of high selectivity and good drug-likeness (QED >0.5, CSS >1.0).

The scatter plot of QED versus CSS (Figure 5B) reveals a positive but weak correlation (Spearman ρ=0.430), suggesting that drug-likeness and selectivity are compatible objectives. Several compounds in Tier 2 and 3 achieve both high selectivity (CSS >1.5) and good drug-likeness (QED >0.5), representing especially attractive candidates for development. The QED versus KEAP1 activity plot (Figure 5C) shows no strong correlation, indicating that potent KEAP1 binders span a range of drug-likeness values.

#### 3.3.3 Lipinski Chemical Space Analysis

Visualization of compounds in Lipinski chemical space [18] (MW vs LogP; TPSA vs LogP) provides insights into the positioning of selective candidates relative to ideal drug space (Figure 8). Most selective compounds occupy the central region of Lipinski space, with molecular weights between 300-550 Da and LogP between 2-6. This positioning is favorable for oral bioavailability.

The MW vs LogP plot (Figure 6A) shows that Tier 2 and 3 compounds cluster within or near the “ideal drug space” (MW <500, LogP 0-5, shaded green). A subset of highly selective compounds fall slightly outside this zone with elevated LogP values (5-7), suggesting that controlled lipophilicity may enhance KEAP1 selectivity. These compounds may still be viable candidates with appropriate formulation strategies or as parenteral therapeutics.

**Figure 6.**
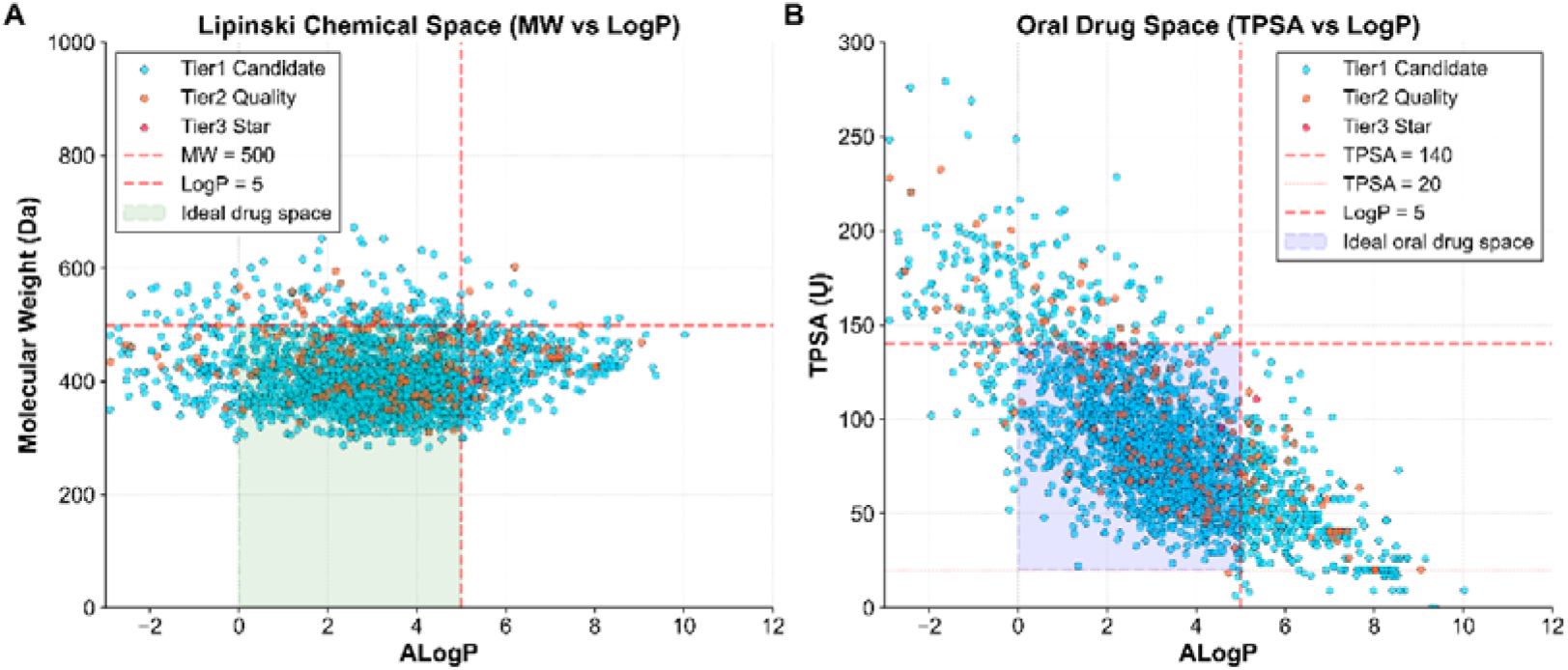
Lipinski and Oral Drug Chemical Space. **(A)** Scatter plot of molecular weight versus ALogP for a random sample of 30,000 compounds, colored by tier. Green shaded region indicates ideal drug space (MW <500, LogP 0-5). Red dashed lines mark Lipinski cutoffs. Most selective candidates (Tier 2 and 3) cluster within or near the ideal space, though some highly selective compounds show elevated LogP. **(B)** Scatter plot of TPSA versus ALogP. Blue shaded region indicates ideal oral drug space (TPSA 20-140, LogP 0-5). The favorable positioning of selective candidates suggests good oral bioavailability potential, though formulation strategies may be needed for the more lipophilic outliers.

The TPSA vs LogP plot (Figure 6B) indicates that most selective candidates occupy the “ideal oral drug space” (TPSA 20-140, LogP 0-5, shaded blue). The favorable TPSA values suggest good permeability and oral absorption potential. Together, these analyses demonstrate that selective Nrf2 activators with acceptable pharmaceutical properties can be identified from natural product libraries, though some trade-offs in lipophilicity may be required for exceptional selectivity.

### 3.4 Chemical Classification and Natural Source Analysis

To understand which chemical classes are enriched among selective candidates and to gain mechanistic insights into the structural basis of selectivity, we performed comprehensive chemical classification analysis.

Tier 2 quality candidates showed striking chemical class enrichment patterns (Figure 7A). The most prominent finding was the strong enrichment of lipids and lipid-like molecules (44.7% of Tier 2 vs 30.0% of all compounds; 1.49-fold enrichment). This enrichment suggests that lipid scaffolds, particularly terpenoids and steroids, may provide favorable frameworks for achieving KEAP1 selectivity. The lipophilic nature of these scaffolds likely facilitates engagement with hydrophobic pockets in the KEAP1 Kelch domain while disfavoring the more promiscuous PXR pocket.

**Figure 7.**
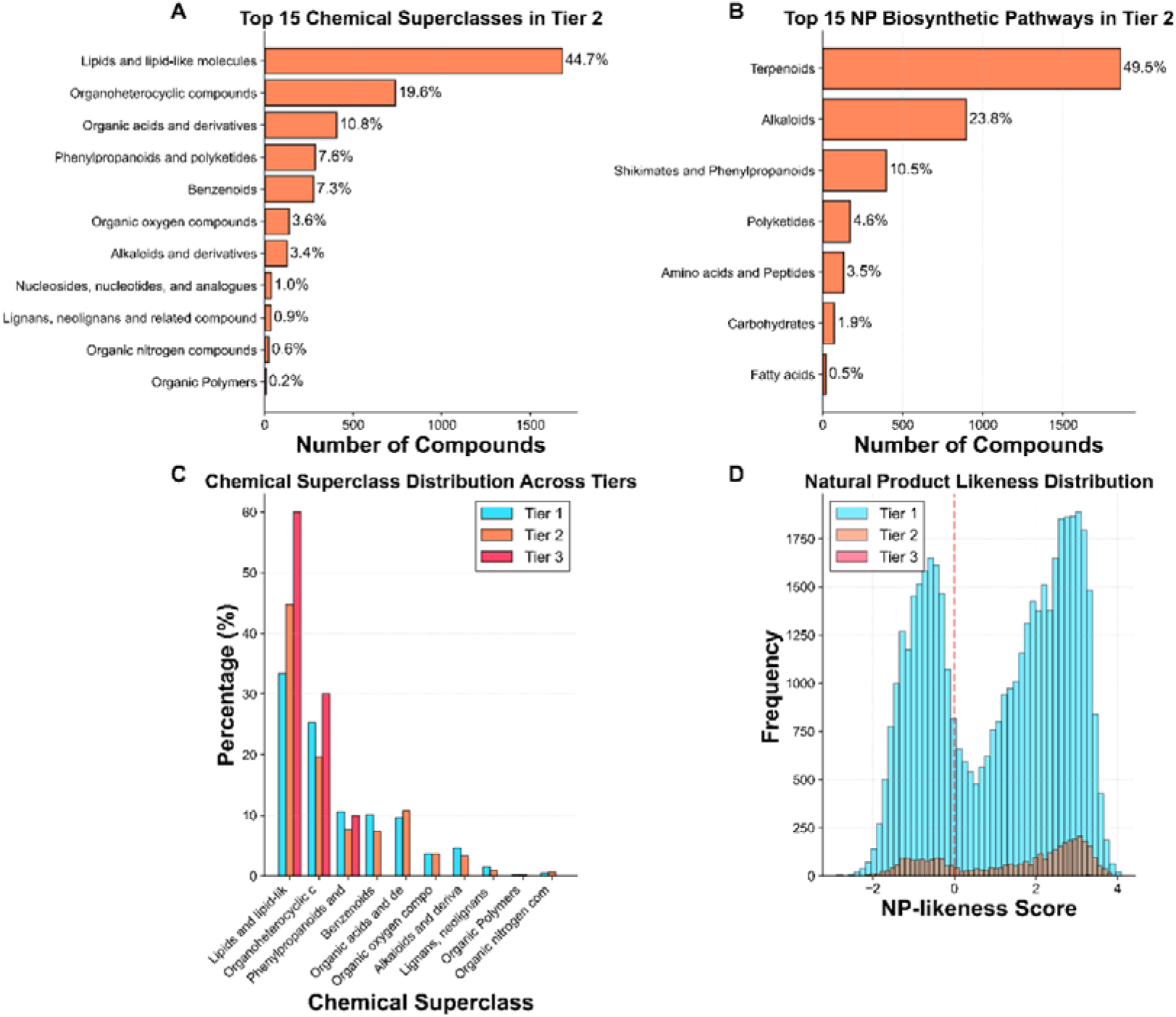
Chemical Classification and Enrichment Analysis. **(A)** Among the high-quality candidates in Tier 2, lipids and lipoid molecules such as terpenes and steroids were significantly enriched (by a factor of 1.49). **(B)** Top 15 NP biosynthetic pathways in Tier 2, revealing diverse origins with terpenoid metabolism predominating. **(C)** Chemical superclass distribution comparison across Tier 1-3. **(D)** NP-likeness score distribution across tiers, showing that selective candidates maintain authentic natural product character. For more details, see Figure S1.

A second remarkable finding was the dramatic enrichment of nucleosides, nucleotides, and analogues (0.96% of Tier 2 vs 0.34% of all compounds; 2.85-fold enrichment). This nearly 3-fold enrichment is particularly intriguing given the relatively small representation of this class in the overall dataset. Nucleoside-based scaffolds may offer unique hydrogen bonding patterns and conformational rigidity that favor KEAP1 binding selectivity.

Conversely, several classes showed depletion among selective candidates. Phenylpropanoids and polyketides (7.6% vs 13.7%; 0.56-fold) and benzenoids (7.4% vs 11.0%; 0.67-fold) were under-represented, suggesting that highly aromatic, planar structures may promote promiscuous binding to multiple targets. [22] This observation aligns with our correlation analysis (Section 3.7) showing that aromatic ring content negatively correlates with selectivity.

Analysis of NP Classifier pathways (Figure 7B) revealed that selective candidates derive from diverse biosynthetic origins. The most common pathways among Tier 2 compounds included terpenoid metabolism, alkaloid biosynthesis, and polyketide synthesis. Notably, compounds from the terpenoid pathway showed particularly favorable selectivity profiles, consistent with the enrichment of lipid-like molecules. This suggests that focusing future screening efforts on terpenoid-rich natural sources (e.g., plants from Lamiaceae family, fungal terpenoids) may yield additional selective candidates.

NP-likeness scores, which quantify how closely a compound resembles known natural products, showed interesting patterns (Figure 7D). Selective candidates maintained positive NP-likeness scores (mean around 0.8-1.2), confirming their authentic natural product character. The preservation of NP-likeness across tiers suggests that selectivity does not require synthetic modifications or departure from natural product space, an important consideration for sourcing and large-scale production

### 3.5 Statistical Correlation and Principal Component Analysis

#### 3.5.1 Correlation Matrix Analysis

To understand the relationships between molecular properties, docking scores, and selectivity indices, we calculated Spearman rank correlations for 17 key variables (Figure 8). Several important patterns emerged from this analysis.

**Figure 8.**
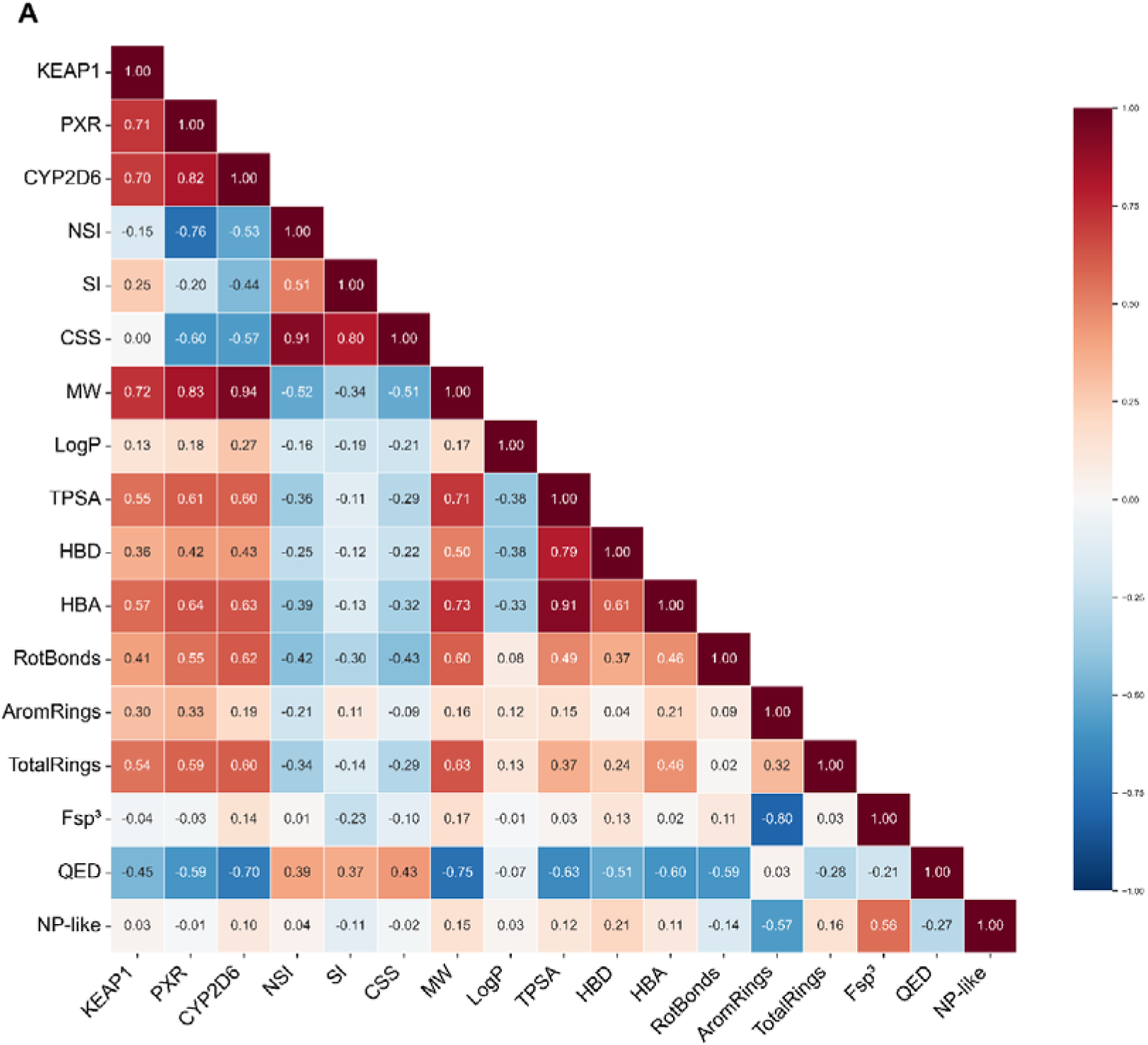
Spearman Correlation Matrix of Molecular Properties and Docking Scores. Lower triangular heatmap showing pairwise correlations between 17 variables including docking scores (KEAP1, PXR, CYP2D6), selectivity indices (NSI, SI, CSS), and molecular properties. Warm colors (red) indicate positive correlations, cool colors (blue) indicate negative correlations.

Selectivity indices showed strong internal correlations. NSI and CSS were highly correlated (ρ=0.907), indicating that compounds with high KEAP1/PXR selectivity also tend to have favorable overall selectivity profiles. SI showed moderate correlation with CSS (ρ=0.798) but weaker correlation with NSI (ρ=0.511), suggesting that CYP2D6 and PXR selectivity are somewhat independent dimensions.

Remarkably, QED showed significant positive correlations with both NSI (ρ=0.391) and CSS (ρ=0.430). This finding is particularly encouraging, as it suggests that drug-like properties and selectivity are compatible and even synergistic objectives. This contrasts with traditional assumptions that achieving high selectivity requires complex, poorly drug-like structures.

Besides that, Aromatic ring count showed negative correlations with NSI (ρ=-0.206) and CSS (ρ=-0.089), supporting our hypothesis that highly aromatic, planar structures promote promiscuous binding. [22] Conversely, Fsp³ (fraction of sp³ carbons) showed weak positive correlation with selectivity, consistent with the principle that threedimensional, non-planar structures enhance selectivity. [21]

KEAP1, PXR, and CYP2D6 scores showed moderate positive correlations (ρ=0.3-0.5), explaining why only 7.38% of compounds passed the Tier 1 criteria despite the 25% expectation for independent scores. Thissuggests that these targets share some promiscuity for certain chemical motifs, making genuine selectivity challenging to achieve.

Moreover, MW correlated moderately with all three target scores (ρ=0.3-0.4), suggesting that larger molecules tend to bind more strongly to all targets. However, MW showed minimal correlation with selectivity indices, indicating that size alone does not determine selectivity—rather, the specific three-dimensional arrangement of functional groups is critical.

The correlation matrix reveals that molecular properties such as aromatic ring count and Fsp3 have measurable impacts on selectivity, providing actionable rules for safety-by-design. The negative correlation between aromatic rings and NSI (ρ=-0.206) suggests that reducing aromaticity can mitigate PXR-related toxicity, a principle that can guide library design. [22] Additionally, the moderate correlations between target scores explain why genuine selectivity is rare, emphasizing the need for multi-parameter optimization in toxicology-aware screening. This analysis underscores that computational tools can decode complex structure-toxicity relationships, enabling proactive risk management.

#### 3.5.2 Principal Component Analysis

PCA on 14 molecular features revealed that the first five principal components explained 78.7% of the total variance (PC1: 27.9%, PC2: 21.1%, PC3: 12.8%, PC4: 10.2%, PC5: 6.8%) (Figure 9A). This indicates substantial redundancy in molecular descriptors, witha few orthogonal dimensions capturing most chemical diversity.

**Figure 9.**
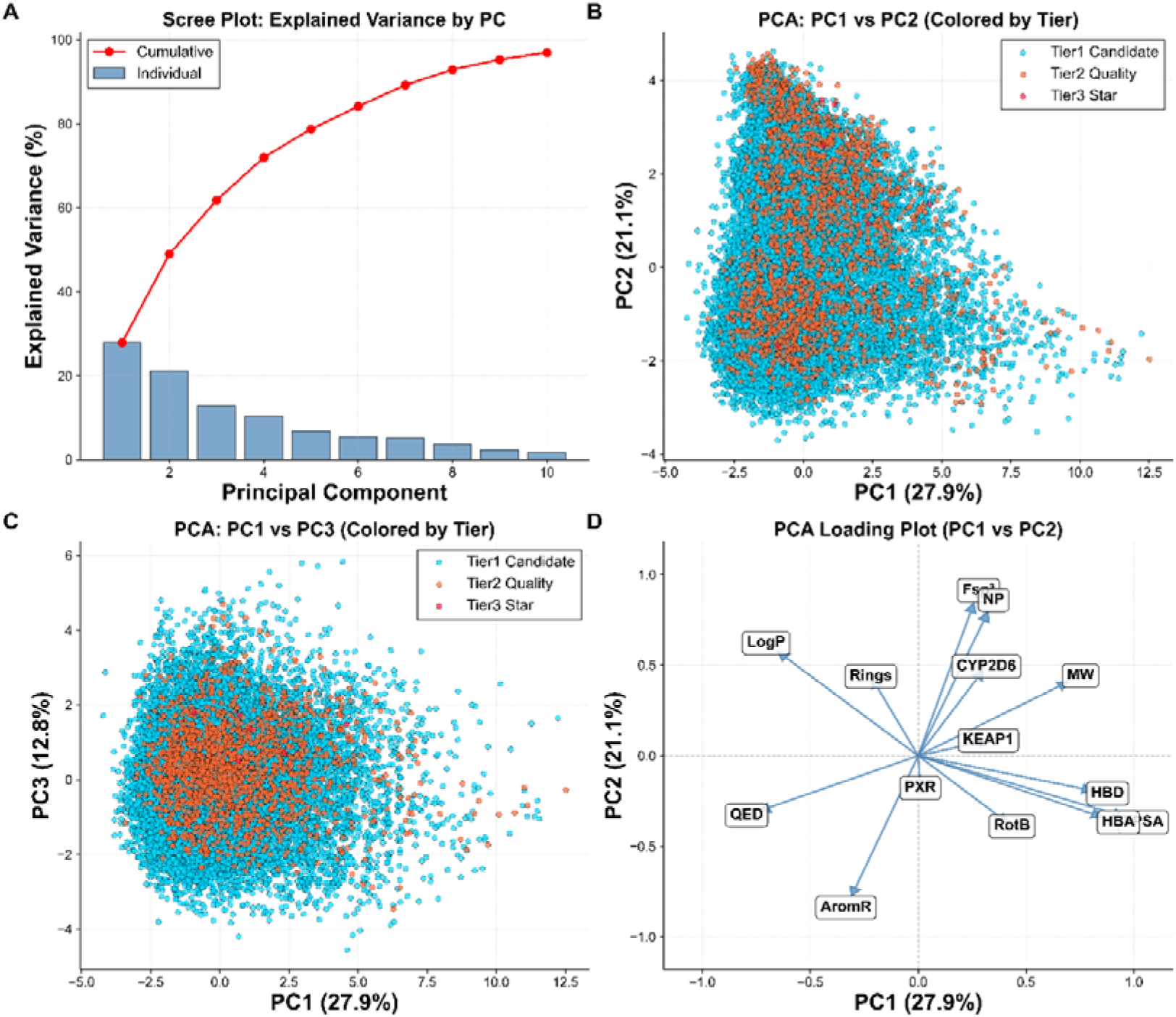
Principal Component Analysis of Molecular Features. **(A)** Scree plot showing explained variance for the first 10 principal components. PC1-PC5 cumulatively explain 78.7% of variance. **(B)** PC1 vs PC2 scatter plot colored by tier, revealing that selective candidates (especially Tier 3) occupy distinct regions of principal component space. **(C)** PC1 vs PC3 scatter plot, providing additional separation along the “drug-likeness” axis. **(D)** Loading plot showing the contribution of each molecular feature to PC1 and PC2. PC1 represents molecular size/complexity, while PC2 represents binding affinity. The positioning of target score vectors explains their intercorrelation and the challenge of achieving selectivity. MW(molecular weight), LogP (ALogP), TPSA, HBD (hydrogen bond donors), HBA (hydrogen bond acceptors), RotB (rotatable bonds), AromR (aromatic rings), Rings (total rings), Fsp³ (fraction sp³), QED, NP (NP-likeness), KEAP1/PXR/CYP2D6 (target scores).

The PC1 vs PC2 scatter plot (Figure 9B) shows that Tier 2 and 3 compounds occupy distinct regions of principal component space compared to the general population. Tier 2 candidates cluster in the positive PC1, positive PC2 quadrant, while Tier 3 compounds show even greater separation. This demonstrates that selective candidates possess coherent combinations of properties that distinguish them from non-selective compounds.

The loading plot (Figure 9D) reveals the molecular properties driving principal component separation. PC1 is dominated by molecular size descriptors (MW, rotatable bonds, HBA), representing a “size/complexity” axis. PC2 is influenced by lipophilicity (LogP) and target scores, representing a “binding affinity” axis. The positioning of KEAP1, PXR, and CYP2D6 vectors in loading space shows that KEAP1 and PXR are somewhat aligned (explaining their correlation), while CYP2D6 is more orthogonal, consistent with our screening strategy of seeking compounds in intermediate CYP2D6 space.

The PC1 vs PC3 plot (Figure 9C) provides additional separation, with PC3 representing a “drug-likeness” axis influenced by QED, aromatic rings, and Fsp³. Selective candidates show favorable positioning along this axis, reinforcing the compatibility of selectivity and drug-likeness.

### 3.6. Tier 3 Star Candidates in Detail

The 10 Tier 3 star candidates represent the most selective compounds identified in this study (Table 2, Figure 10). These compounds achieved the most stringent screening criteria, with KEAP1 scores >90th percentile, PXR scores <10th percentile, and CYP2D6 scores in the narrow 40-60th percentile window. Four of these compounds ranked in the top 20 by CSS.

**Figure 10.**
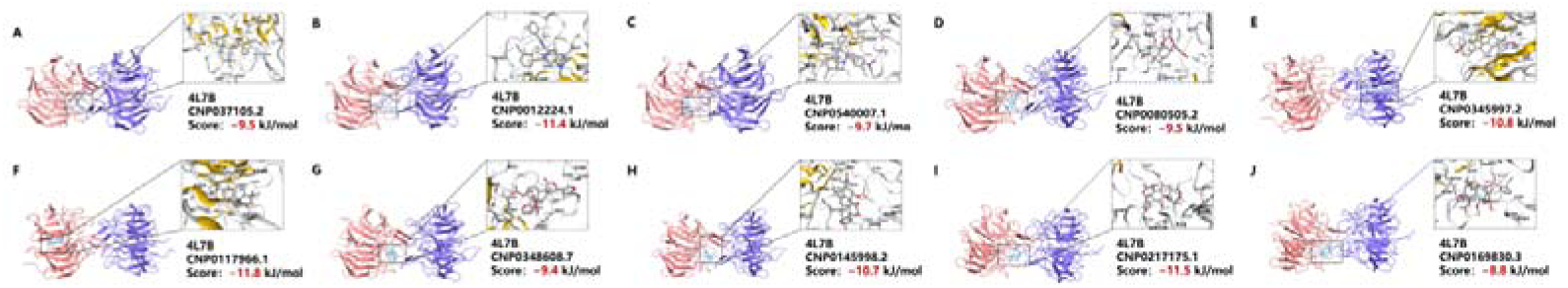
Molecular docking diagrams of the top ten complexes with the 4L7B protein.

The #1 ranked compound CNP0371205.2 (CSS=2.64, NSI=12.29) exemplifies the ideal selectivity profile: strong KEAP1 binding (20.91), minimal PXR binding (1.60), moderate CYP2D6 interaction (36.87), and favorable drug-likeness (MW: 377.5 Da, QED: 0.52). This compound belongs to the phenylpropanoid class and represents a novel scaffold for Nrf2 activation. CNP0012224.1 (CSS=2.02, NSI=6.38), a complex organoheterocyclic compound featuring imidazopyridine and oxadiazole rings, demonstrates that selectivity can also be achieved with nitrogen-rich heterocycles. CNP0540007.1 (CSS=1.87, NSI=5.92), a benzodiazepine derivative with nitrophenyl substituents, represents an unexpected scaffold for selective KEAP1 engagement.

Among the remaining Tier 3 compounds, several lipid-based structures stand out. CNP0117966.1 (CSS=1.62), CNP0348608.7 (CSS=1.52), and CNP0145998.2 (CSS=1.50) all feature complex polycyclic terpenoid cores with diverse oxygenation patterns.

These compounds likely engage KEAP1 through a combination of hydrophobic interactions with the Kelch domain groove and specific hydrogen bonds with key residues.

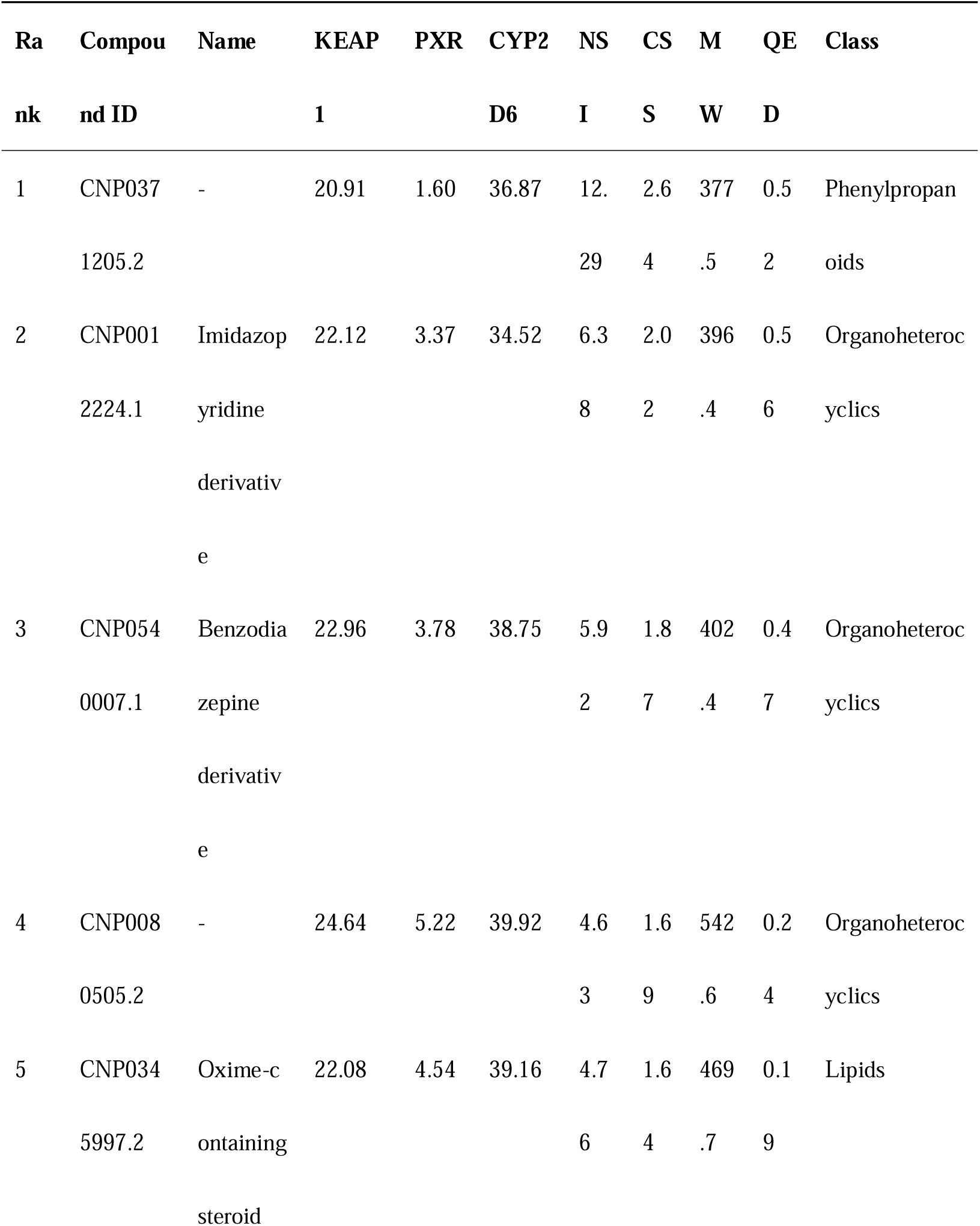
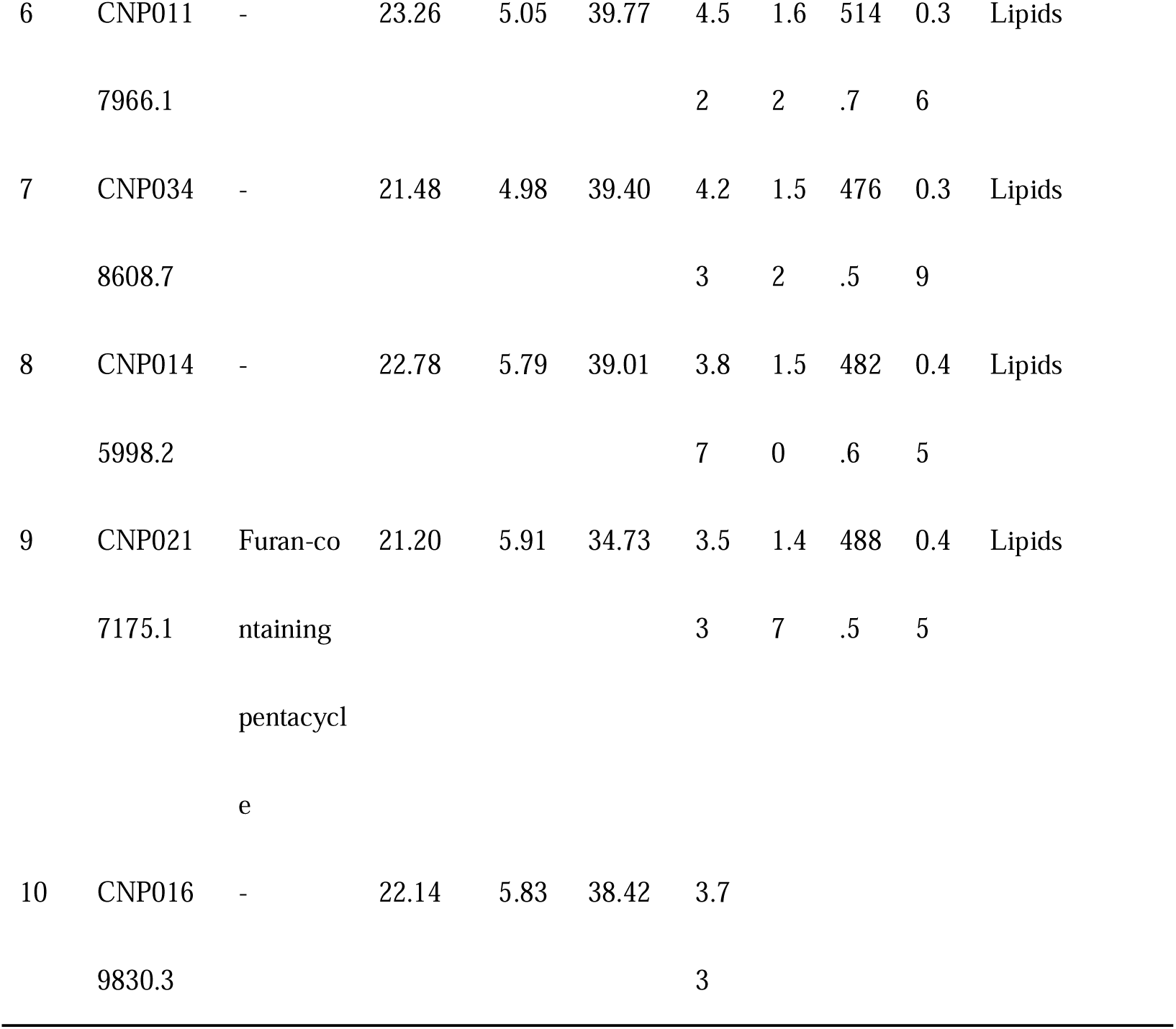

**Figure 10. Molecular docking diagrams of the top ten complexes with the 4L7B protein.**
| Ra | Compou | Name | KEAP | PXR | CYP2 | NS | CS | M | QE | Class |
| --- | --- | --- | --- | --- | --- | --- | --- | --- | --- | --- |
| nk | nd ID |  | 1 |  | D6 | I | S | W | D |  |
| 1 | CNP037 | - | 20.91 | 1.60 | 36.87 | 12. | 2.6 | 377 | 0.5 | Phenylpropan |
|  | 1205.2 |  |  |  |  | 29 | 4 | .5 | 2 | oids |
| 2 | CNP001 | Imidazop | 22.12 | 3.37 | 34.52 | 6.3 | 2.0 | 396 | 0.5 | Organoheteroc |
|  | 2224.1 | yridine |  |  |  | 8 | 2 | .4 | 6 | yclics |
|  |  | derivativ |  |  |  |  |  |  |  |  |
|  |  | e |  |  |  |  |  |  |  |  |
| 3 | CNP054 | Benzodia | 22.96 | 3.78 | 38.75 | 5.9 | 1.8 | 402 | 0.4 | Organoheteroc |
|  | 0007.1 | zepine |  |  |  | 2 | 7 | .4 | 7 | yclics |
|  |  | derivativ |  |  |  |  |  |  |  |  |
|  |  | e |  |  |  |  |  |  |  |  |
| 4 | CNP008 | - | 24.64 | 5.22 | 39.92 | 4.6 | 1.6 | 542 | 0.2 | Organoheteroc |
|  | 0505.2 |  |  |  |  | 3 | 9 | .6 | 4 | yclics |
| 5 | CNP034 | Oxime-c | 22.08 | 4.54 | 39.16 | 4.7 | 1.6 | 469 | 0.1 | Lipids |
|  | 5997.2 | ontaining |  |  |  | 6 | 4 | .7 | 9 |  |
|  |  | steroid |  |  |  |  |  |  |  |  |
| 6 | CNP011 | - | 23.26 | 5.05 | 39.77 | 4.5 | 1.6 | 514 | 0.3 | Lipids |
|  | 7966.1 |  |  |  |  | 2 | 2 | .7 | 6 |  |
| 7 | CNP034 | - | 21.48 | 4.98 | 39.40 | 4.2 | 1.5 | 476 | 0.3 | Lipids |
|  | 8608.7 |  |  |  |  | 3 | 2 | .5 | 9 |  |
| 8 | CNP014 | - | 22.78 | 5.79 | 39.01 | 3.8 | 1.5 | 482 | 0.4 | Lipids |
|  | 5998.2 |  |  |  |  | 7 | 0 | .6 | 5 |  |
| 9 | CNP021 | Furan-co | 21.20 | 5.91 | 34.73 | 3.5 | 1.4 | 488 | 0.4 | Lipids |
|  | 7175.1 | ntaining |  |  |  | 3 | 7 | .5 | 5 |  |
|  |  | pentacycl |  |  |  |  |  |  |  |  |
|  |  | e |  |  |  |  |  |  |  |  |
| 10 | CNP016 | - | 22.14 | 5.83 | 38.42 | 3.7 |  |  |  |  |
|  | 9830.3 |  |  |  |  | 3 |  |  |  |  |

### 3.7 Candidate Prioritization and Property Profiling

To identify the most promising candidates for further development, we prioritized compounds from both Tier 2 and Tier 3 pools based on a combined assessment of selectivity and drug-likeness. Figure 16 presents a comprehensive property comparison of the top 20 candidates ranked by CSS and filtered by QED ≥ 0.3, revealing important insights into the relationship between selectivity metrics and physicochemical properties.

Notably, the final Top 20 cohort comprises 4 Tier 3 (ultra-selective) and 16 Tier 2 (high-quality) compounds, rather than exclusively Tier 3 candidates. This composition reflects our prioritization strategy that balances exceptional selectivity with favorable drug-like properties. Six Tier 3 compounds (CNP0080505.2, CNP0345997.2, CNP0348608.7, CNP0145998.2, CNP0217175.1, and CNP0169830.3) were excluded due to QED values below 0.3, indicating suboptimal physicochemical profiles despite their high selectivity. Importantly, two Tier 2 compounds (CNP0361771.4, CSS = 3.31; CNP0118918.2, CSS = 3.23) demonstrated superior CSS values compared to most Tier 3 candidates, suggesting that exceptional selectivity can be achieved while maintaining robust drug-likeness.

The property comparison across nine key metrics reveals several critical patterns (Figure 11). The progressive increase in CSS from bottom to top (Figure 11A) validates the ranking criterion and highlights the stratification of selectivity performance. Panels B and C (NSI and SI) show consistent trends with CSS, confirming that compounds with high comprehensive selectivity also exhibit strong Nrf2-specific selectivity and safety profiles. The docking scores (Figure 11D–11F) demonstrate an inverse relationship between PXR scores (Figure 11E) and overall selectivity, emphasizing the importance of off-target minimization in achieving high CSS values. This observation is particularly relevant given PXR’s role in drug-drug interactions and metabolic regulation.

**Figure 11.**
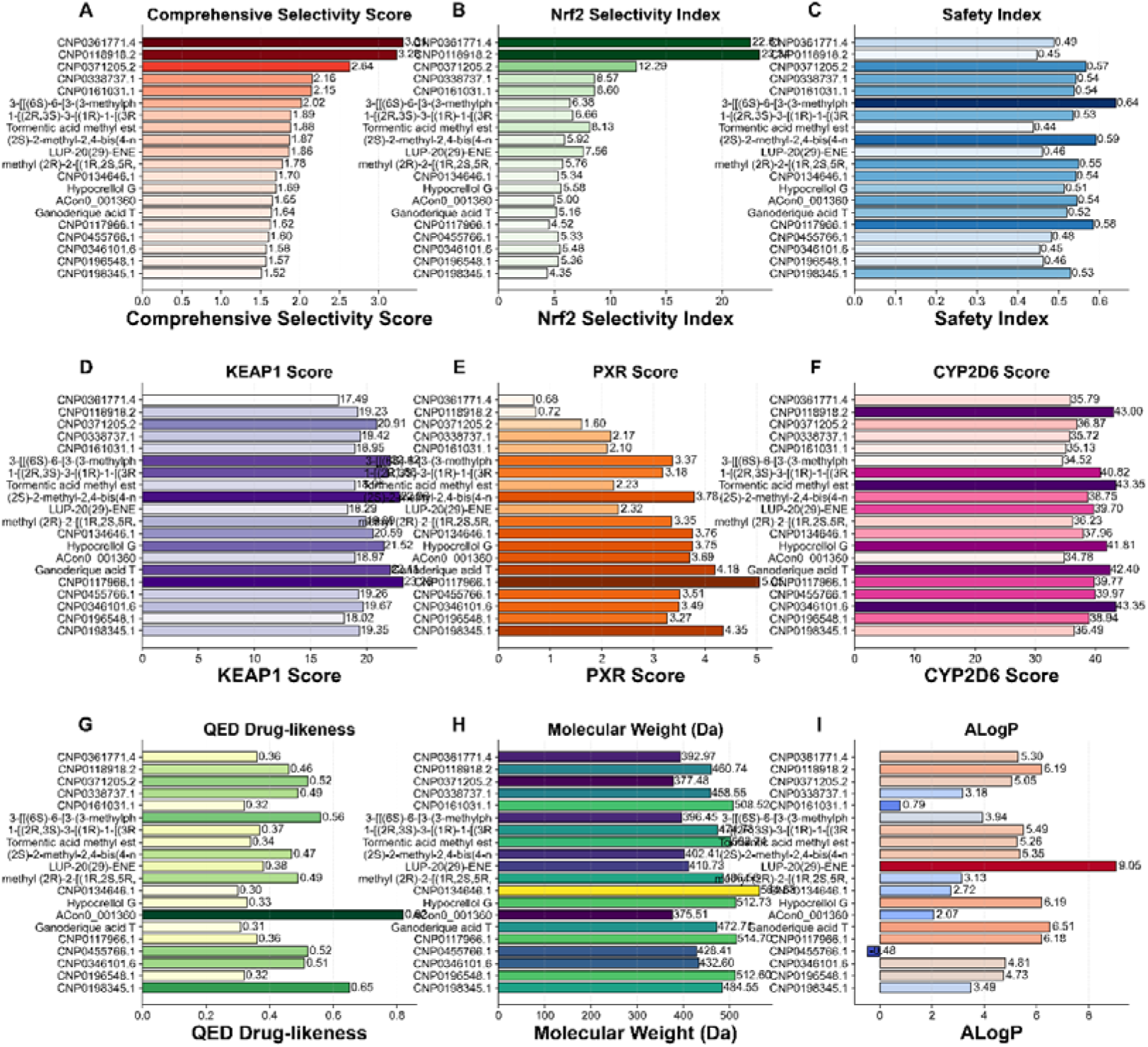
Top 20 Candidates: Property Comparison. Horizontal bar charts comparing the top 20 candidates (sorted by CSS, lowest to highest) across nine key properties: (A) CSS, (B) NSI, (C) SI, (D) KEAP1 score, (E) PXR score (note: lower is better), (F) CYP2D6 score, (G) QED, (H) Molecular weight, and (I) ALogP. Bars are color-coded by normalized property values. The progressive increase in CSS from bottom to top demonstrates the ranking criterion. Note the inverse relationship between PXR scores and overall selectivity. Candidates were selected from combined Tier 2 and Tier 3 pools, ranked by CSS (descending), and filtered by QED ≥ 0.3 to ensure drug development potential. The resulting Top 20 comprises 4 Tier 3 (ultra-selective) and 16 Tier 2 (high-quality) compounds, reflecting an optimal balance between selectivity and drug-likeness. Six Tier 3 compounds with QED < 0.3 were excluded due to suboptimal physicochemical properties.

The drug-likeness metrics (Figure 11G–11I) reveal that all Top 20 candidates maintain favorable properties, with QED values ranging from 0.30 to 0.82, molecular weights within acceptable limits, and ALogP values suggesting reasonable membrane permeability. The color-coded visualization using normalized colormaps facilitates rapid identification of property outliers and confirms the overall homogeneity of the top candidate cohort.

## 4. Conclusion

This work establishes a safety-by-design framework for discovering selective Nrf2 activators by integrating KEAP1, PXR, and CYP2D6 into the largest natural-product docking campaign conducted to date. From 628,898 molecules, we identified ultraselective candidates with strong KEAP1 affinity and minimized off-target risks, demonstrating the value of simultaneous efficacy–toxicity screening. Beyond providing promising lead scaffolds, this study contributes a three-target molecular docking dataset and a complete, scalable pipeline that can guide future drug design and predictive toxicology efforts.

## Acknowledgments

Special thanks to Dr. Yuxuan Lyv for their invaluable advice and Ms. Nature Belle for her assistance. We also extend our appreciation to Aging laboratory technicians for their diligent work and to the administrative staff for their support throughout the project. Furthermore, we would like to express our gratitude to UESTC_BioMed for their revisions of the figures and language in this manuscript.

## Ethics approval and consent to participate

Not applicable.

## Consent for publication

Not applicable.

## Author Contributions

Conceptualization, Y.W..; methodology, Y.W., H.M., R.L. and L.F.; software, H.C. and R.L; validation, Y.W.; formal analysis, H.M..; investigation, H.M.; resources, Y.W.; data curation, Y.W.; writing—original draft preparation, H.M.; writing—review and editing, Y.W., H.M., H.C..; visualization, L.F.. and Y.W.; supervision, H.M.; project administration, H.M.; All authors have read and agreed to the published version of the manuscript.

---

**Figure S1.**
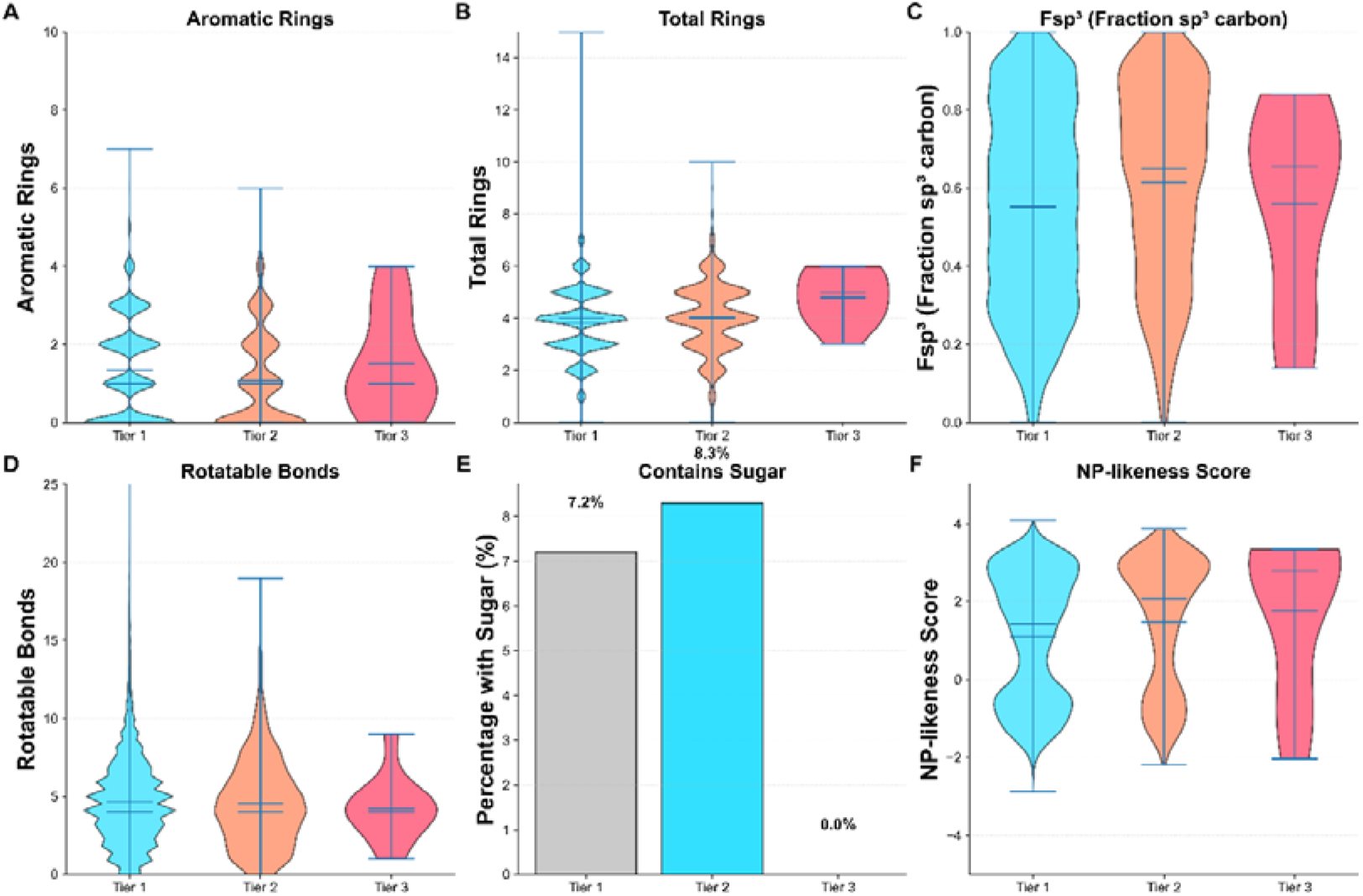
Structural Features Analysis Across Tiers. Distribution of key structural features: (A) Aromatic ring count, (B) Total ring count, (C) Fraction of sp³ carbons (Fsp³), (D) Rotatable bonds, (E) Sugar content (percentage), and (F) NP-likeness. Lower aromatic ring content and higher Fsp³ values in selective candidates suggest that three-dimensional, non-planar structures favor selectivity. The moderate sugar content (∼15-20%) indicates that glycosylation is compatible with but not required for selectivity. These structural patterns provide design principles for scaffold optimization and library design.

